# SCAPP: An algorithm for improved plasmid assembly in metagenomes

**DOI:** 10.1101/2020.01.12.903252

**Authors:** David Pellow, Alvah Zorea, Maraike Probst, Ori Furman, Arik Segal, Itzhak Mizrahi, Ron Shamir

## Abstract

**Background:** Metagenomic sequencing has led to the identification and assembly of many new bacterial genome sequences. These bacteria often contain plasmids: usually small, circular double-stranded DNA molecules that may transfer across bacterial species and confer antibiotic resistance. These plasmids are generally less studied and understood than their bacterial hosts. Part of the reason for this is insufficient computational tools enabling the analysis of plasmids in metagenomic samples.

**Results:** We developed SCAPP (Sequence Contents-Aware Plasmid Peeler) - an algorithm and tool to assemble plasmid sequences from metagenomic sequencing. SCAPP builds on some key ideas from the Recycler algorithm while improving plasmid assemblies by integrating biological knowledge about plasmids.

We compared the performance of SCAPP to Recycler and metaplasmidSPAdes on simulated metagenomes, real human gut microbiome samples, and a human gut plasmidome dataset that we generated. We also created plasmidome and metagenome data from the same cow rumen sample and used the parallel sequencing data to create a novel assessment procedure. Overall, SCAPP outperformed Recycler and metaplasmidSPAdes across this wide range of datasets.

**Conclusions:** SCAPP is an easy to use Python package that enables the assembly of full plasmid sequences from metagenomic samples. It outperformed existing metagenomic plasmid assemblers in most cases, and assembled novel and clinically relevant plasmids in samples we generated such as a human gut plasmidome. SCAPP is open-source software available from: https://github.com/Shamir-Lab/SCAPP.

## Background

Plasmids play a critical role in microbial adaptation, such as antibiotic resistance or other metabolic capabilities, and genome diversification through horizontal gene transfer. However, plasmid evolution and ecology across different microbial environments and populations are poorly characterized and understood. Thousands of plasmids have been sequenced and assembled directly from isolated bacteria, but constructing complete plasmid sequences from short read data remains a hard challenge. The task of assembling plasmid sequences from shotgun metagenomic sequences, which is our goal here, is even more daunting.

There are several reasons for the difficulty of plasmid assembly. First, plasmids represent a very small fraction of the sample’s DNA and thus may not be fully covered by the read data in high-throughput sequencing experiments. Second, they often share sequences with the bacterial genomes and with other plasmids, resulting in tangled assembly graphs. For these reasons, plasmids assembled from bacterial isolates are usually incomplete, fragmented into multiple contigs, and contaminated with sequences from other sources. The challenge is reflected in the title of a recent review on the topic: “On the (im)possibility of reconstructing plasmids from whole-genome short-read sequencing data” [1]. In a metagenomic sample, these problems are amplified since the assembly graphs are much larger, more tangled and fragmented.

There are a number of tools that can be used to detect plasmid sequences including PlasmidFinder [2], cBar [3], gPlas [4], PlasFlow [5], and others. There is also the plasmidSPAdes assembler for assembling plasmids in isolate samples [6]. However, there are currently only two tools that attempt to reconstruct complete plasmid sequences in metagenomic samples: Recycler [7] and metaplasmidSPAdes [8] (mpSpades). mpSpades iteratively generates smaller and smaller subgraphs of the assembly graph by removing contigs with coverage below a threshold that increases in each iteration. As lower coverage segments of the graph are removed, longer contigs may be constructed in the remaining subgraph. Cyclic contigs are considered as putative plasmids and then verified using the profile of their genetic contents. The main idea behind Recycler is that a single shortest circular path through each node in the assembly graph can be found efficiently. The circular paths that have uniform read coverage are iteratively “peeled” off the graph and reported as possible plasmids. The peeling process reduces the residual coverage of each involved node, or removes it altogether. We note that these tools, as well as our work, focus on circular plasmids and do not assemble linear plasmid sequences.

Here we present SCAPP (Sequence Contents-Aware Plasmid Peeler), a new algorithm that uses the peeling idea of Recycler and also leverages external biological knowledge about plasmid sequences. In SCAPP the assembly graph is annotated with plasmid-specific genes (PSGs) and nodes are assigned weights reflecting the chance that they are plasmidic based on a plasmid sequence classifier [9]. In the annotated assembly graph we prioritize peeling off circular paths that include plasmid genes and highly probable plasmid sequences. SCAPP also uses the PSGs and plasmid scores to filter out likely false positives from the set of potential plasmids.

We tested SCAPP on both simulated and diverse real metagenomic data and compared its performance to Recycler and mpSpades. Overall, SCAPP performed better than the other tools across these datasets. SCAPP has higher precision than Recycler in all cases, meaning it more accurately constructs correct plasmids from the sequencing data. SCAPP also has higher recall than mpSpades in most cases, and higher precision in most of the real datasets. We developed and tested a novel strategy given parallel plasmidome and metagenome sequencing of the same sample. We show how to accurately assess the performance of the tools on metagenome data, even in the absence of known reference plasmids.

## Implementation

SCAPP accepts as input a metagenomic assembly graph, with nodes representing the sequences of assembled contigs and edges representing *k*-long sequence overlaps between contigs, and the paired-end reads from which the graph was assembled. SCAPP processes each component of the assembly graph and iteratively assembles plasmids from them. The output of SCAPP is a set of cyclic sequences representing confident plasmid assemblies.

A high-level overview of SCAPP is provided in Box 1 and depicted graphically in Figure 1; the full algorithmic details are presented below. For brevity, we describe only default parameters below, see Supplementary Information, section S1 for alternatives.

**Figure 1.**
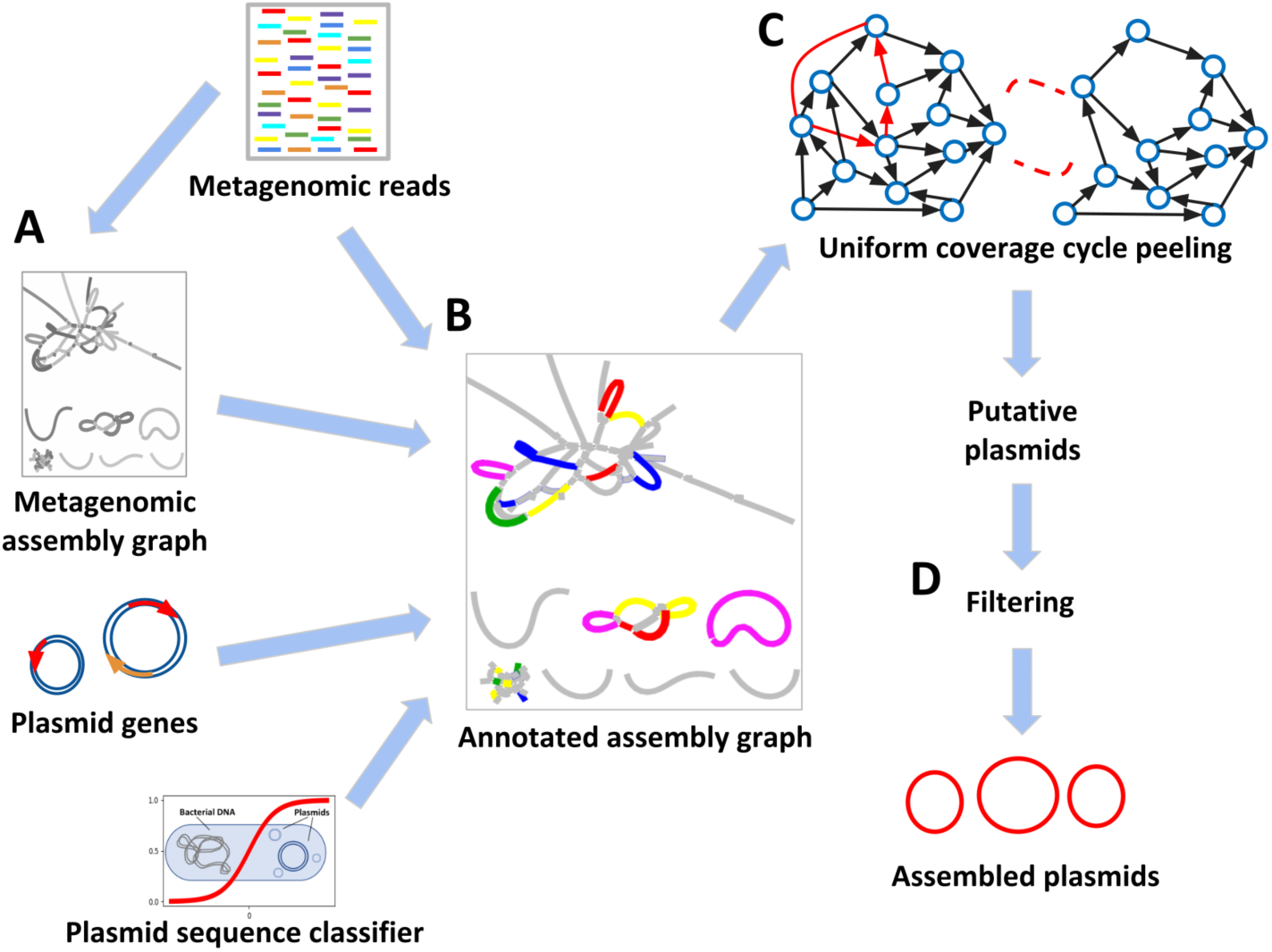
Graphical overview of the SCAPP algorithm. A: The metagenomic assembly graph is created from the sample reads. B: The assembly graph is annotated with read mappings, presence of plasmid specific genes, and node weights based on sequence length, coverage, and plasmid classifier score. C: Potential plasmids are iteratively peeled from the assembly graph. An efficient algorithm finds cyclic paths in the annotated assembly graph that have low weight and high chance of being plasmids. Cycles with uniform coverage are peeled. D: Confident plasmid predictions are retained using plasmid sequence classification and plasmid-specific genes to remove likely false positive potential plasmids.

SCAPP is available from https://github.com/Shamir-Lab/SCAPP, and fully documented there. It was written in Python3 and can be installed as a conda package, directly from Bioconda or from its sources.

### The SCAPP algorithm

The full SCAPP algorithm is given in Algorithm 1. The peel function, which defines how cycles are peeled from the graph, is given in Algorithm 2.

### Read mapping

The first step in creating the annotated assembly graph (Box 1 step 1a) is to align the reads to the contigs in the graph. The links between paired-end reads aligning across contig junctions are used to evaluate potential plasmid paths in the graph. SCAPP performs read alignment using BWA [10] and the alignments are filtered to retain only primary read mappings, sorted, and indexed using SAMtools [11].

#### Algorithm 1 SCAPP pipeline

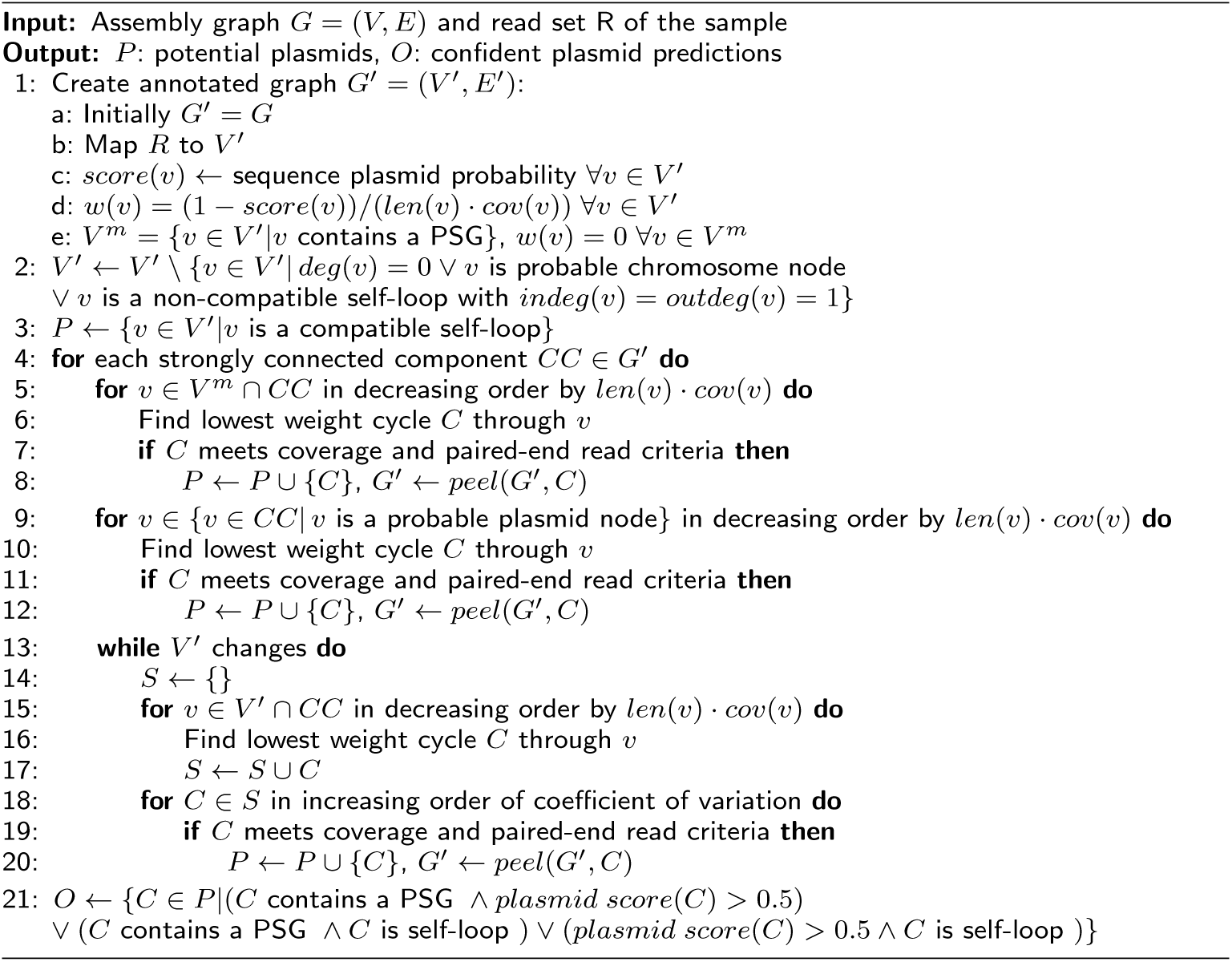

#### Algorithm 2 peel(G, C)

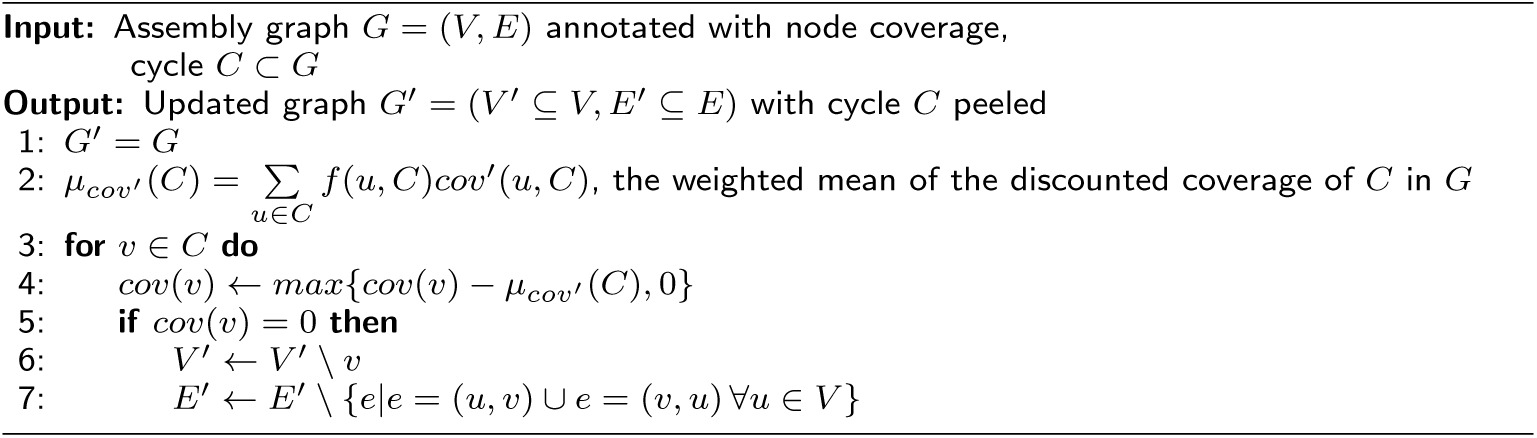

#### Box 1.

**Overview of SCAPP**

1. Annotate the assembly graph:
  a. Map reads to nodes of the assembly graph
  b. Find nodes with plasmid-specific gene matches
  c. Compute plasmid sequence scores of nodes
  d. Assign node weights
2. **for** each strongly connected component **do**
3. Iteratively peel uniform coverage cycles through plasmid gene nodes
4. Iteratively peel uniform coverage cycles through high scoring nodes
5. Iteratively peel shortest cycle through each remaining node if it meets plasmid criteria
6. Output the set of confident plasmid predictions

### Plasmid-specific gene annotation

We created sets of PSGs by database mining and curation by plasmid microbiology experts from the Mizrahi Lab (Ben-Gurion University). Information about these PSG sets is found in Supplementary Information, section S2. The sequences them- selves are available from https://github.com/Shamir-Lab/SCAPP/scapp/data.

A node in the assembly graph is annotated as containing a PSG hit (Box 1 step 1b) if there is a BLAST match between one of the PSG sequences and the sequence corresponding to the node (≥ 75% sequence identity along ≥ 75% of the length of the gene).

### Plasmid sequence score annotation

We use PlasClass [9] to annotate each node in the assembly graph with a plasmid score (Box 1 step 1c). PlasClass uses a set of logistic regression classifiers for sequences of different lengths to assign a classification score reflecting the likelihood of each node to be of plasmid origin.

We re-weight the node scores according to the sequence length as follows. For a given sequence of length *L* and plasmid probability *p* assigned by the classifier, the re-weighted plasmid score is: 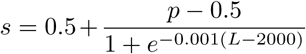. This tends to pull scores towards 0.5 for short sequences, for which there is lower confidence, while leaving scores of longer sequences practically unchanged.

Long nodes (*L* > 10 kbp) with low plasmid score (*s <* 0.2) are considered probable chromosomal sequences and are removed, simplifying the assembly graph. Similarly, long nodes (*L* > 10 kbp) with high plasmid score (*s* > 0.9) are considered probable plasmid nodes.

### Assigning node weights

In order to apply the peeling idea, nodes are assigned weights (Box 1 step 1d) so that lower weights correspond to higher likelihood to be assembled into a plasmid. Plasmid score and PSG annotations are incorporated into the node weights. A node with plasmid score *s* is assigned a weight *w*(*v*) = (1−*s*)*/*(*C·L*) where *C* is the depth of coverage of the node’s sequence and *L* is the sequence length. This gives lower weight to nodes with higher coverage, longer sequence and higher plasmid scores. Nodes with PSG hits are assigned a weight of zero, making them more likely to be integrated into any lowest-weight cycle in the graph that can pass through them.

### Finding low-weight cycles in the graph

The core of the SCAPP algorithm is to iteratively find a lowest weight (“lightest”) cycle going through each node in the graph for consideration as a potential plasmid. We use the bidirectional single-source, single-target shortest path implementation of the NetworkX Python package [12].

The order that nodes are considered matters since in each iteration potential plasmids are peeled from the graph, affecting the cycles that may be found in subsequent iterations. The plasmid annotations are used to decide the order that nodes are considered: first all nodes with PSGs, then all probable plasmid nodes, and then all other nodes in the graph (Box 1 step 2). If the lightest cycle going through a node meets certain criteria described below, it is peeled off, changing the coverage of nodes in the graph. Performing the search for light cycles in this order ensures that the cycles through more likely plasmid nodes will be considered before other cycles.

### Assessing coverage uniformity

The lightest cyclic path, weighted as described above, going through each node is found and evaluated. Recycler sought a cycle with near uniform coverage, reasoning that all contigs that form a plasmid should have roughly the same coverage. However, this did not take into account the overlap of the cycle with other paths in the graph (see Figure 2). To account for this, we instead compute a discounted coverage score for each node in the cycle based on its interaction with other paths as follows:

**Figure 2.**
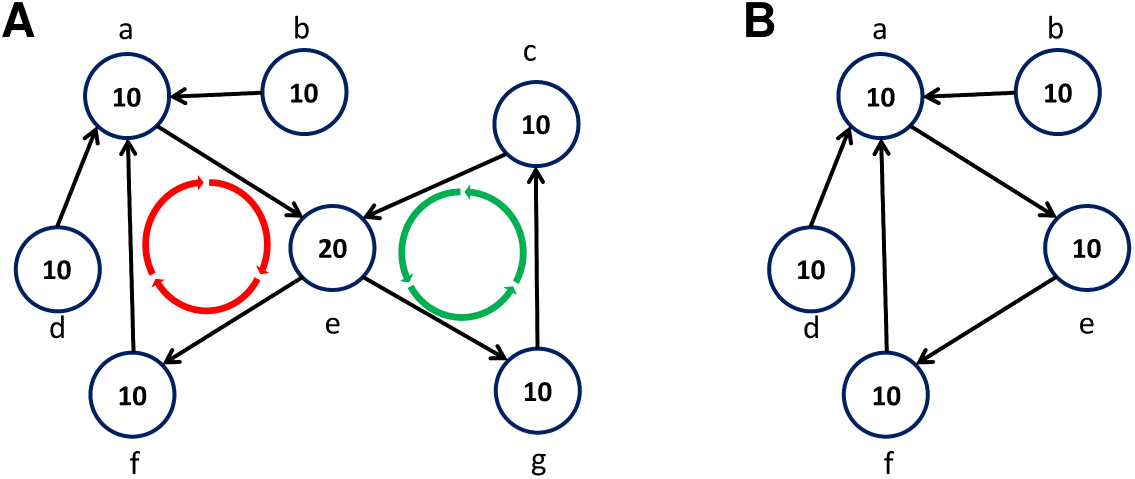
Evaluating and peeling cycles. Numbers inside nodes indicate coverage. All nodes in the example have equal length. A: Cycles (*a, e, f*) and (*c, e, g*) have the same average coverage (13.33) and coefficient of variation (CV, 0.35), but their discounted CV values differ: The discounted coverage of node *a* is 6, and the discounted coverage of node *e* is 10 in both cycles. The left cycle has discounted CV=0.22 and the right has discounted CV=0. By peeling off the mean discounted coverage of the right cycle (10) one gets the graph in B. Note that nodes *g, c* were removed from the graph since their coverage was reduced to 0, and the coverage of node *e* was reduced to 10.

The *discounted coverage* of a node *v* in the cycle *C* is its coverage *cov*(*v*) times the fraction of the coverage on all its neighbors (both incoming and outgoing), 𝒩 (*v*), that is on those neighbors that are in the cycle (see Figure 2):

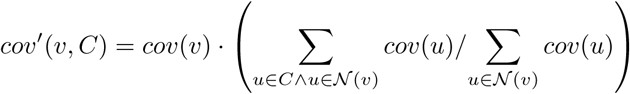

A node *v* in cycle *C* with contig length *len*(*v*) is assigned a weight *f* corresponding to its fraction of the length of the cycle: 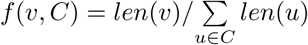. These weights are used to compute the weighted mean and standard deviation of the discounted coverage of the nodes in the cycle: 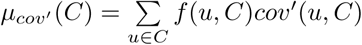,

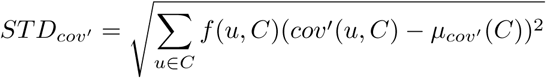

The coefficient of variation of *C*, which evaluates its coverage uniformity, is the ratio of the standard deviation to the mean:

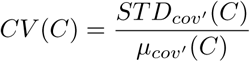

### Finding potential plasmid cycles

After each lightest cycle has been generated, it is evaluated as a potential plasmid based on its structure in the assembly graph, the PSGs it contains, its plasmid score, paired-end read links, and coverage uniformity. The precise evaluation criteria are described in Supplementary Information, section S3. A cycle that passes them is defined as a potential plasmid (Box 1 steps 3-5). The potential plasmid cycles are peeled from the graph in each iteration as defined in Algorithm 2 (see also Figure 2).

### Filtering confident plasmid assemblies

In the final stage of SCAPP, PSGs and plasmid scores are used to filter out likely false positive plasmids from the output and create a set of confident plasmid assemblies (Box 1 step 6). All potential plasmids are assigned a length-weighted plasmid score and are annotated with PSGs as was done for the contigs during graph annotation. Those that belong to at least two of the following sets are reported as confident plasmids: (a) potential plasmids containing a match to a PSG; (b) potential plasmids with plasmid score > 0.5; (c) self-loop nodes.

## Results

We tested SCAPP on simulated metagenomes, human gut metagenomes, a human gut plasmidome dataset that we generated and also on parallel metagenome and plasmidome datasets from the same cow rumen microbiome specimen that we generated. The test settings and evaluation methods are described in Supplementary Information, section S5.

### Simulated metagenomes

We created seven read datasets simulating metagenomic communities of bacteria and plasmids and assembled them. Datasets of increasing complexity were created as shown in Table 1. We randomly selected bacterial genomes along with their associated plasmids, and used realistic distributions for genome abundance and plasmid copy number. Further details of the simulation can be found in Supplementary Information, section S4. 5M paired-end reads were generated for Sim1 and Sim2, 10M for Sim3 and Sim4, and 20M for Sim5, Sim6, and Sim7.

**Table 1.** Performance on simulated metagenome datasets. The number of covered plasmids (# covered) reports the number of the simulation plasmids that were covered by reads along at least 95% of their length. The set of covered plasmids is used as the gold standard in calculating the performance metrics. The numbers in parentheses are the median plasmid lengths (in kbp). F1 score is presented as a percent.

| Sample | #<br>genomes | #<br>plasmids | #<br>covered | Recycler |  | mpSpades |  | SCAPP |  |
| --- | --- | --- | --- | --- | --- | --- | --- | --- | --- |
|  |  |  |  | #<br>plasmids | F1 | #<br>plasmids | F1 | #<br>plasmids | F1 |
| Sim1 | 10 | 9 (5.9) | 9 | 7 (5.6) | 50.0 | 1 (4.3) | 20.0 | 5 (5.6) | 57.1 |
| Sim2 | 50 | 47 (19.3) | 37 | 20 (3.8) | 40.1 | 9 (5.0) | 39.1 | 23 (5.5) | 43.3 |
| Sim3 | 200 | 210 (22.4) | 136 | 61 (3.6) | 32.8 | 27 (7.0) | 32.3 | 48 (5.8) | 42.9 |
| Sim4 | 200 | 177 (25.4) | 132 | 62 (4.1) | 40.8 | 29 (6.0) | 36.5 | 51 (6.2) | 48.9 |
| Sim5 | 300 | 318 (23.9) | 253 | 115 (3.6) | 35.2 | 53 (5.1) | 33.8 | 100 (6.5) | 47.5 |
| Sim6 | 400 | 480 (13.5) | 368 | 138 (3.0) | 28.5 | 59 (5.5) | 27.1 | 118 (5.5) | 36.5 |
| Sim7 | 500 | 571 (17.3) | 410 | 132 (3.5) | 31.1 | 69 (5.3) | 28.1 | 141 (5.2) | 40.5 |

Table 1 presents features of the simulated datasets and reports the performance of Recycler, mpSpades, and SCAPP on them. For brevity we report only F1 scores; precision and recall scores are reported in Supplementary Table 1, Supplementary Information (section S6). SCAPP had the highest F1 score in all cases, followed by Recycler. SCAPP consistently achieved higher precision than Recycler, allowing it to perform better overall. mpSpades had the highest precision, but assembled far fewer plasmids than the other tools and gained lower recall and F1 scores. This suggests that mpSpades mostly assembled the easier plasmids. Indeed, most of the plasmids assembled by mpSpades were also assembled by the other tools (see Figure S1 in Supplementary Information).

### Human gut microbiomes

We tested the plasmid assembly algorithms on data of twenty publicly available human gut microbiome samples selected from the study of Vrieze *et al*. [13]. The true set of plasmids in these samples is unknown. Instead, we matched all assembled contigs to PLSDB [14] and considered the set of the database plasmids that were covered by the contigs as the gold standard (see Supplementary Information, section S5 for details). All tools were evaluated according to the same gold standard. We note that this limits the evaluation to known plasmids. We chose the human gut microbiome in this experiment and the next, as it is one of the most widely studied microbiome environments so plasmids in gut microbiome samples are most likely to be represented in the database.

Table 2 presents the results of the three algorithms averaged across all twenty samples. The detailed results on each of the samples are presented in Supplementary Table 2 and Figure S2, Supplementary Information (section S7). SCAPP performed best in more cases, with mpSpades failing to assemble any gold standard plasmid in over half the samples. We note that all of the cases where SCAPP had recall of 0 occurred when the number of gold standard plasmids was very small and the other tools also failed to assemble them. On the largest samples with the most gold standard plasmids SCAPP performed best, highlighting its superior performance on the types of samples most likely to be of interest in experiments aimed at plasmid assembly. SCAPP consistently outperformed Recycler by achieving higher precision, a result that is consistent with the other experiments.

**Table 2.**
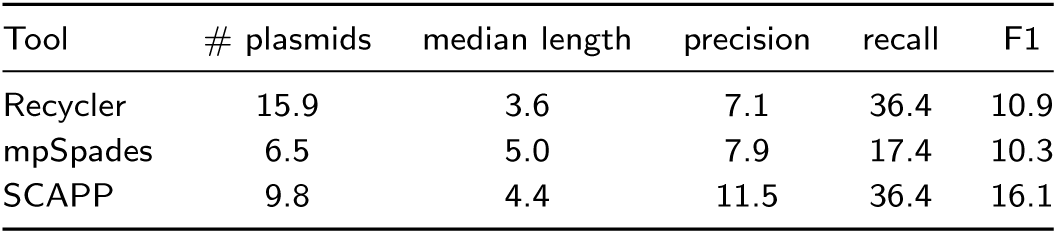
Performance on the human gut metagenomes. Number of plasmids, the median plasmid length (in kbp), and performance measures for all tools are averaged across the twenty samples. The average number of plasmids and median length of the gold standard sets of plasmids were 4.8 and 12.4 respectively

### Human gut plasmidome

The protocol developed in Brown Kav *et al*. [15] enables extraction of DNA from isolate or metagenomic samples with the plasmid content highly enriched. The sequence contents of such a sample is called the *plasmidome* of the sample. This enrichment for plasmid sequences increases the chance of revealing the plasmids in the sample. The protocol was assessed to achieve samples with at least 65% plasmid contents by Krawczyk *et al*. [5]. We sequenced the plasmidome of the human gut microbiome from a healthy adult male according to the plasmid enrichment protocol. 18,616,649 paired-end reads were sequenced with the Illumina HiSeq2000 platform, read length 150bp and insert size 1000.

The gold standard set of plasmids, determined as for the gut metagenome samples, consisted of 74 plasmids (median length = 2.1 kbp). Performance was computed as in the metagenomic samples and is shown in Table 3. SCAPP achieved best overall performance, while mpSpades had lower precision and much lower recall than the other tools.

**Table 3.** Performance on the human gut plasmidome. Number of plasmids, the median plasmid length (in kbp), and performance measures for all tools.

| Tool | # plasmids | median length | precision | recall | F1 |
| --- | --- | --- | --- | --- | --- |
| Recycler | 93 | 2.1 | 15.1 | 37.8 | 21.5 |
| mpSpades | 53 | 3.0 | 11.3 | 9.4 | 10.3 |
| SCAPP | 82 | 2.4 | 17.1 | 35.9 | 23.1 |

Notably, although the sample was obtained from a healthy donor, some of the plasmids reconstructed by SCAPP matched reference plasmids found in potentially pathogenic hosts such as *Klebsiella pneumoniae*, pathogenic serovars of *Salmonella enterica*, and *Shigella sonnei*. The detection of plasmids previously isolated from pathogenic hosts in the healthy gut indicates potential pathways for transfer of virulence genes.

We used MetaGeneMark [16] to find potential genes in the plasmids assembled by SCAPP. 294 genes were found, and we annotated them with the NCBI nonredundant (nr) protein database using BLAST. 46 of the plasmids contained 170 (58%) genes with matches in the database, of which 77 (45%) had known functional annotations, which we grouped manually in Figure 3A. There were six antibiotic and toxin resistance genes, all on plasmids that were not in the gold standard set, highlighting SCAPP’s ability to find novel resistance carrying plasmids. 60 of the 77 genes (78%) with functional annotations had plasmid associated functions: replication, mobilization, recombination, resistance, and toxin-antitoxin systems. 29 out of the 33 plasmids that contained functionally annotated genes (88%) contained at least one of these plasmid associated functions. This provides a strong indication that SCAPP succeeded in assembling true plasmids of the human gut plasmidome.

**Figure 3.**
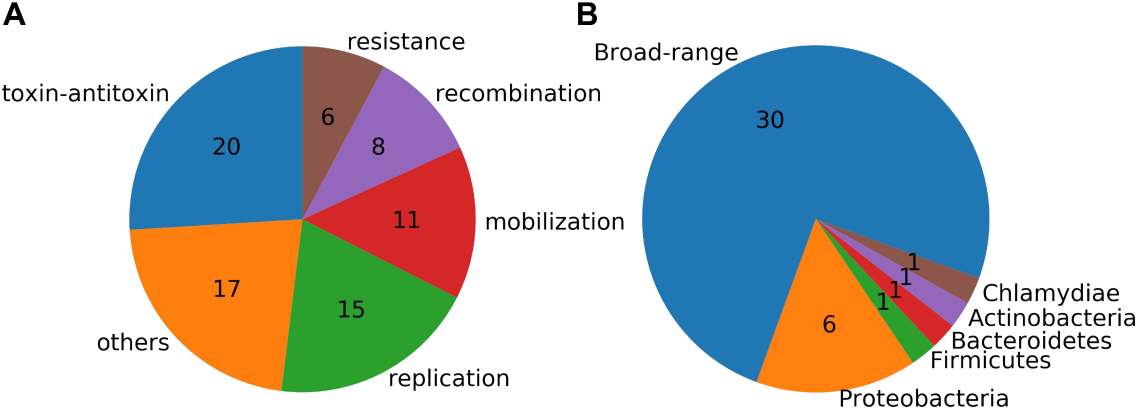
Annotation of genes on the plasmids identified by SCAPP in the human gut plasmidome sample. A: Functional annotations of the plasmid genes. B: Host annotations of the plasmid genes. “Broad-range” plasmids had genes annotated with hosts from more than one phylum.

We also examined the hosts that were annotated for the plasmid genes and found that almost all of the plasmids with annotated genes contained genes with annotations from a variety of hosts, which we refer to here as “broad-range” (see Figure 3B). Of the 40 plasmids with genes from annotated hosts, only 10 (25%) had genes with annotated hosts all within a single phylum. This demonstrates that these plasmids assembled and identified by SCAPP may be involved in one stage of transferring genes, such as the antibiotic resistance genes we detected, across a range of bacteria.

### Parallel metagenomic and plasmidome samples

We performed two sequencing assays on the same cow rumen microbiome sample of a four month old calf. In one subsample total DNA was sequenced. In the other, plasmid-enriched DNA was extracted as described in Brown Kav *et al*. [15] and sequenced (see Figure 4). 27,127,784 paired-end reads were sequenced in the plasmidome, and 54,292,256 in the metagenome. Both were sequenced on the Illumina HiSeq2000 platform with read length 150bp and insert size 1000.

**Figure 4.**
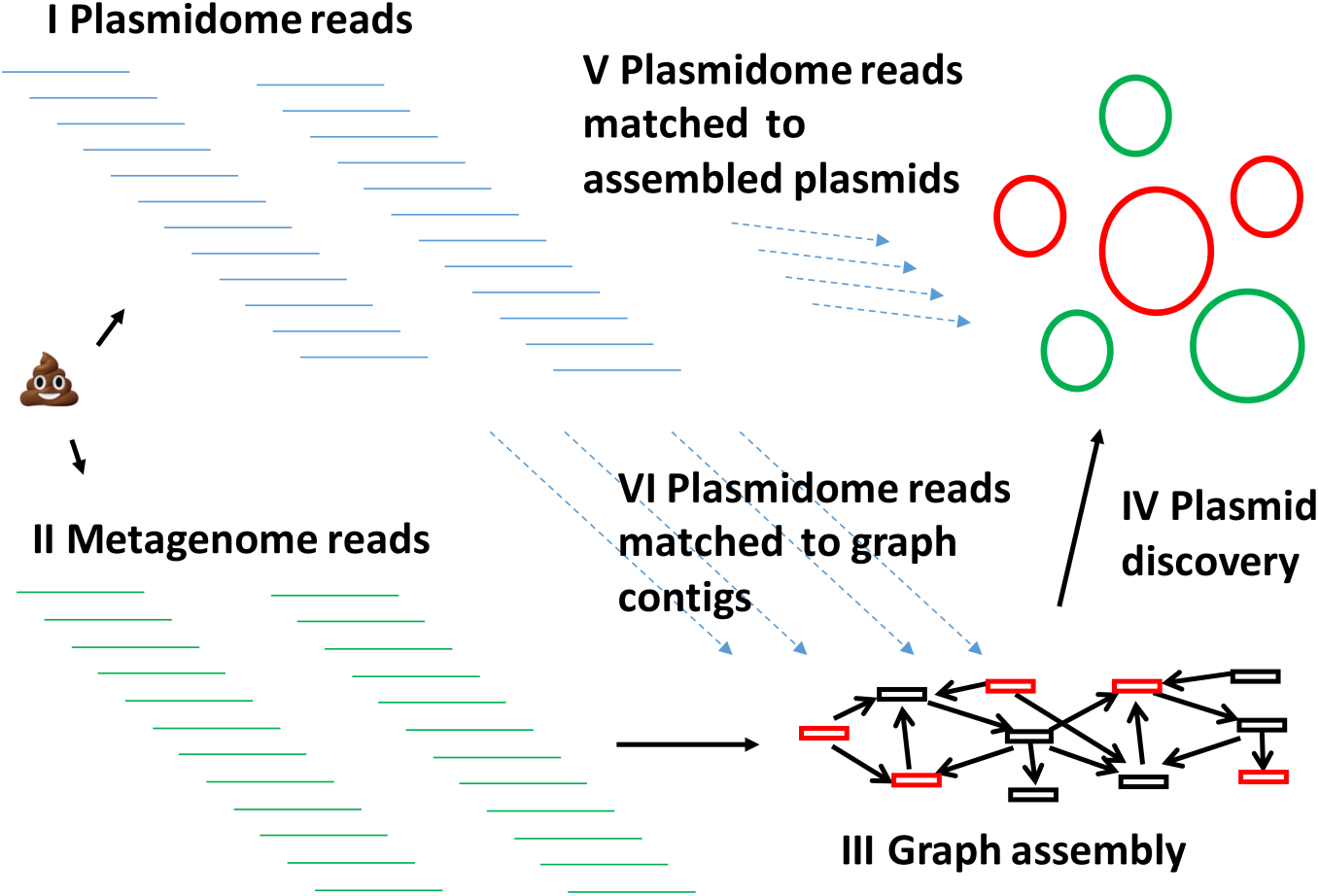
Outline of the read-based performance assessment. Plasmidome (I) and metagenome reads (II) are obtained from subsamples of the same sample. III: The metagenome reads are assembled into a graph. IV: The graph is used to detect and report plasmids by the algorithm of choice. V: The plasmidome reads are matched to assembled plasmids. Matched plasmids (red) are used to calculate plasmid read-based precision. VI: The plasmidome reads are matched to the assembly graph contigs. Covered contigs (red) are considered plasmidic. The fraction of total length of plasmidic contigs included in the detected plasmids gives the plasmidome read-based recall.

This parallel data enabled us to assess the plasmids assembled on the metagenome using the plasmidome, without resorting to PLSDB matches as the gold standard. Such assessment is especially useful for samples from non-clinical environments such as the cow rumen, as PLSDB likely under-represents plasmids in them.

Table 4 summarizes the results of the three plasmid discovery algorithms on both subsamples. mpSpades made the fewest predictions and Recycler made the most. To compare the plasmids identified by the different tools, we considered two plasmids to be the same if their sequences matched at > 80% identity across > 90% of their length. The comparison is shown in Figure S3, Supplementary Information (section S8). In the plasmidome subsample, 50 plasmids were identified by all three methods. Seventeen were common to the three methods in the metagenome. In both subsamples, the Recycler plasmids included all or almost all of those identified by the other methods plus a large number of additional plasmids. In the plasmidome, SCAPP and Recycler shared many more plasmids than mpSpades and Recycler.

**Table 4.** Number of plasmids assembled by each tool and their median lengths (in kbp) for the parallel metagenome and plasmidome samples.

| Tool | metagenome |  | plasmidome |  |
| --- | --- | --- | --- | --- |
|  | # plasmids | median length | # plasmids | median length |
| Recycler | 60 | 4.3 | 147 | 1.7 |
| SCAPP | 25 | 5.8 | 110 | 1.8 |
| mpSpades | 26 | 6.2 | 65 | 2.0 |

We also evaluated the results of the plasmidome and metagenome assemblies by comparison to PLSDB as was done for the human gut samples. The metagenome contained only one matching PLSDB reference plasmid, and none of the tools assembled it. The plasmidome had only seven PLSDB matches, and mpSpades, Recycler, and SCAPP had F1 scores of 2.86, 2.67, and 1.74, respectively. The low fraction of PLSDB matches out of the assembled plasmids suggests that the tools can identify novel plasmids that are not in the database.

In order to fully leverage the power of parallel samples, we computed the performance of each tool on the metagenomic sample using the reads of the plasmidomic sample, without doing any contig and plasmid assembly on the latter. The rationale was that the reads of the plasmidome represent the full richness of plasmids in the sample in a way that is not biased by a computational procedure or prior biological knowledge.

We calculated the *plasmidome read-based precision* by mapping the plasmidomic reads to the plasmids assembled from the metagenomic sample (Figure 4). A plasmid with > 90% of its length covered by more than one plasmidomic read was considered to be a true positive. The precision of an algorithm was defined as the fraction of true positive plasmids out of all reported plasmids. The *plasmidome read-based recall* was computed by mapping the plasmidomic reads to the contigs of the metagenomic assembly. Contigs with > 90% of their length covered by plasmidomic reads at depth > 1 were called *plasmidic contigs*. Plasmidic contigs that were part of the assembled plasmids were counted as true positives, and those that were not were considered false negatives. The recall was defined as the fraction of the plasmidic contigs’ length that was integrated in the assembled plasmids. Note that the precision and recall here are measured using different units (plasmids and base pairs, respectively) so they are not directly related. For mpSpades, which does not output a metagenomic assembly, we mapped the contigs from the metaSPAdes assembly to the mpSpades plasmids using BLAST (> 80% sequence identity matches along > 90% of the length of the contigs).

There were 293 plasmidic contigs in the metagenome assembly graph, with a total length of 146.6 kbp. The plasmidome read-based performance is presented in Figure 5A. All tools achieved a similar recall of around 12. SCAPP and mpSpades performed similarly, with SCAPP having slightly higher precision (24.0 vs 23.1) but slightly lower recall (11.9 vs 12.2). Recycler had a bit higher recall (13.1), but at the cost of far lower precision (11.7). Hence, a much lower fraction of the plasmids assembled by Recycler in the metagenome were actually supported by the parallel plasmidome sample, adding to the other evidence that the false positive rate of Recycler exceeds that of the other tools.

**Figure 5.**
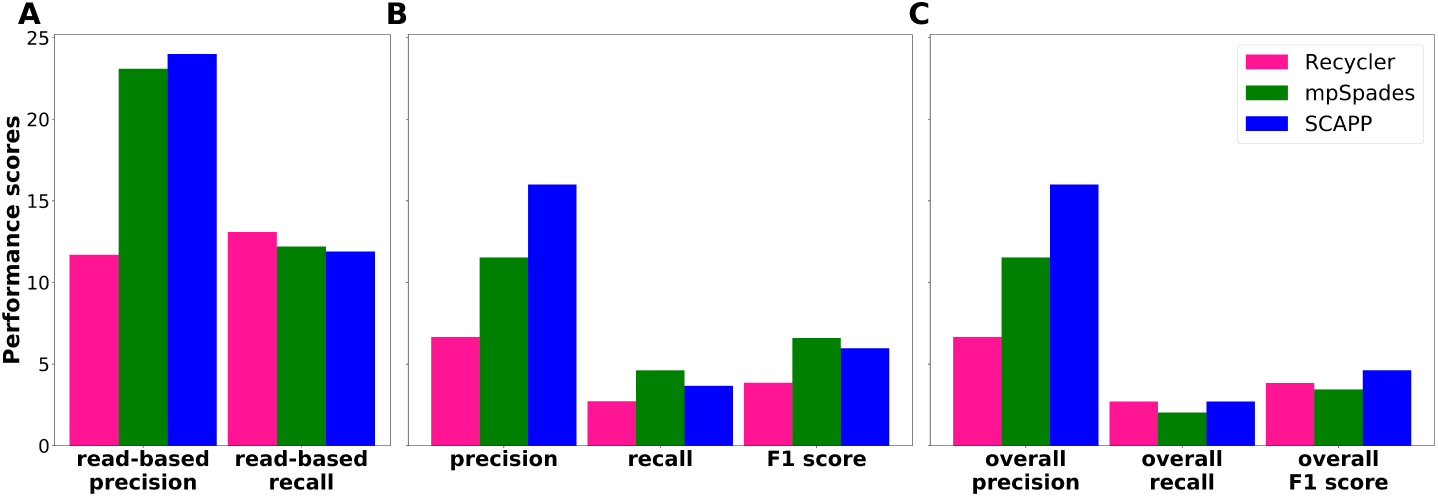
Performance on the parallel datasets. A: Plasmidome read-based performance. B: Performance of each tool on the plasmids assembled from the metagenome using as gold standard the plasmids assembled from the plasmidome by the same tool. C: Overall performance on the plasmids assembled from the metagenome compared to the union of all plasmids assembled by all tools in the plasmidome.

We also compared the plasmids assembled by each tool in the two subsamples. For each tool, we considered the plasmids it assembled from the plasmidome to be the gold standard set, and used it to score the plasmids it assembled in the metagenome. The results are shown in Figure 5B. SCAPP had the highest precision. Since mpSpades had a much smaller gold standard set, it achieved higher recall and F1. Recycler output many more plasmids than the other tools in both samples, but had much lower precision, suggesting that many of its plasmid predictions may be spurious.

Next, we considered the union of the plasmids assembled across all tools as the gold standard set and recomputed the scores. We refer to them as “overall” scores. Figure 5C shows that overall precision scores were the same as in Figure 5B, while overall recall was lower for all the tools, as expected. mpSpades underperformed because of its smaller set of plasmids, and SCAPP had the highest overall F1 score.

We detected potential genes in the plasmids assembled by SCAPP in the plasmidome sample and annotated them as we did for the human gut plasmidome. The gene function and host annotations are shown in Figure S4, Supplementary Information (section S8). Out of 242 genes, only 34 genes from 17 of the plasmids had annotations, and only 18 of these had known functions, highlighting that many of the plasmids in the cow rumen plasmidome are as yet unknown. The high percentage of genes of plasmid function (15/18) indicates that SCAPP succeeded in assembling novel plasmids. Unlike in the human plasmidome, most of the plasmids with known host annotations had hosts from a single phylum.

### Performance summary

We summarize the performance of the tools across all the test datasets in Table 5. The performance of two tools was considered similar (denoted ≈) if their scores were within 5% of each other. Performance of one tool was considered to be much higher than the other (≫) if its score was > 30% higher (an increase of 5 − 30% is denoted by >).

**Table 5.** Summary of performance. Comparison of the performance of the tools on each of the datasets. When multiple samples were tested, the number of samples appears in parentheses, and average performance is reported. For the parallel samples results are for the evaluation of the metagenome based on the plasmidome, and precision and recall are plasmidome read-based. Unless otherwise stated, F1 score is used. Note that in the simulations, SCAPP ≫ mpSpades.

| Test | Ranking |
| --- | --- |
| Simulations (7) | SCAPP > Recycler > mpSpades |
| Human gut metagenomes (20) | SCAPP ≫ mpSpades > Recycler |
| Plasmidome | SCAPP > Recycler ≫ mpSpades |
| Parallel: within tool | mpSpades > SCAPP ≫ Recycler |
| Parallel: “overall”, across tools | SCAPP > Recycler > mpSpades |
| Parallel: precision | SCAPP ≈ mpSpades ≫ Recycler |
| Parallel: recall | Recycler > mpSpades ≈ SCAPP |

We see that in most cases SCAPP was the highest performer. Furthermore, in all other cases SCAPP performed close to the top performing tool.

### Resource usage

The runtime and memory usage of the three tools are presented in Table 6. Recycler and SCAPP require assembly by metaSPAdes and pre-processing of the reads and the resulting assembly graph. SCAPP also requires post-processing of the assembled plasmids. mpSpades requires post-processing of the assembled plasmids with the plasmidVerify tool. The reported runtimes are for the full pipelines necessary to run each tool – from reads to assembled plasmids.

**Table 6.** Resource usage of the three methods. Peak RAM of the assembly step (metaSPAdes for Recycler and SCAPP, metaplasmidSPAdes for mpSpades) in GB. Runtime (wall clock time, in minutes) is reported for the entire pipeline including assembly and any pre-processing and post-processing required. Human metagenome results are an average across the 20 samples.

| Dataset | RAM (GB) | Runtime (minutes) |  |  |
| --- | --- | --- | --- | --- |
|  |  | Recycler | mpSpades | SCAPP |
| Human metagenomes | 21 | 115 | 103 | 130 |
| Plasmidome | 30 | 907 | 548 | 909 |
| Parallel metagenome | 148 | 2118 | 2132 | 2230 |
| Parallel plasmidome | 26 | 881 | 684 | 884 |

In almost all cases assembly was the most memory intensive step, and so all tools achieved very similar peak memory usage (within 0.01 GB). Therefore, we report the RAM usage for this step.

The assembly step was also the longest step in all cases. SCAPP was slightly slower than Recycler as a result of the additional annotation steps, and mpSpades was 5 – 40% faster. However, note that mpSpades does not output a metagenomic assembly graph, so users interested in both the plasmid and non-plasmid sequences in a sample would need to run metaSPAdes as well, practically doubling the runtime.

Performance measurements were made on a 44-core, 2.2 GHz server with 792 GB of RAM. 16 processes were used where possible. Recycler is single-threaded, so only one process was used for it.

## Discussion

Plasmid assembly from metagenomic sequencing is a very difficult task, akin to finding needles in a haystack. This difficulty is demonstrated by the low numbers of plasmids found in real samples. Even in samples of the human gut microbiome, which is widely studied, relatively few plasmids that have matches in the extensive plasmid database PLSDB were recovered. Despite the challenges, SCAPP was able to assemble plasmids across a number of clinically relevant samples. SCAPP significantly outperformed mpSpades in simulation and on a range of human gut metagenome and plasmidome samples. In simulation mpSpades achieved very high precision at the expense of low recall, and SCAPP had higher combined F1 score. The high precision was not observed in real data, which is more difficult than the simulations. SCAPP was also consistently better than Recycler across almost all tests. Though SCAPP and Recycler share the idea of cycle peeling, SCAPP was shown to have higher precision, due to incorporating additional biological information and better edge weighting.

Another contribution of this study is the joint analysis of the parallel metagenome and plasmidome from the same sample. We show that this enables a novel way to evaluate plasmid assembly algorithms on the metagenome data, by using the coverage information from the plasmidome. This novel approach bypasses the need to rely on known plasmids for evaluation, which is biased due to research focus. We developed several evaluation metrics for such data, and think they can be useful for future plasmid studies, especially in non-clinical and non-human samples where plasmid knowledge is scarce.

A key difficulty in evaluation of performance of plasmid discovery algorithms is the lack of gold standard. The verification of reported plasmids is done either based on prior biological knowledge, which is biased, or by experimental verification, which is slow and expensive. Moreover, such verification evaluates precision but does not give information on the extent of missed plasmids, or recall. While simulations can evaluate both parameters accurately, they are inherently artificial, and necessitate many modeling assumptions that are not fully supported by experimental data. For that reason we chose here to focus primarily on real data, and preferred diversity in the real data types over extensive but artificial simulations. The parallel samples strategy is another partial answer to this problem.

SCAPP has several limitations. Like the other de Bruijn graph-based plasmid assemblers, it may split a cycle into two when a shorter cycle is a sub-path of a longer cycle. It also has difficulties in finding very long plasmids, as these tend to not be completely covered and fragmented into many contigs in the graph. Note however that it produced longer cycles than Recycler. Compared to mpSpades, each algorithm produced longer cycles in different tests. Another limitation is the inherent bias in relying on known plasmid genes and plasmid databases, which tend to under-represent non-clinical samples. With further use of tools like SCAPP, perhaps with databases tailored to specific environments, further improvement is possible.

## Conclusions

We introduced SCAPP, a new plasmid discovery tool based on combination of graph theoretical and biological considerations. Overall, SCAPP demonstrated better performance than Recycler and metaplasmidSpades in a wide range of real samples from diverse contexts. By applying SCAPP across large sets of samples, many new plasmid reference sequences can be assembled, enhancing our understanding of plasmid biology and ecology.

## Supplementary information for

## S1 Alternatives for user set parameters

The SCAPP pipeline is highly flexible, and many of the options and parameters can be set by the user. In most cases, we recommend using the default options and settings. Some of the alternatives that can be chosen by the user are described below. All of the parameter settings that may be changed by the user are fully documented at: https://github.com/Shamir-Lab/SCAPP.

### Read mapping

The user has the option of providing a sorted and indexed BAM alignment file created by any method.

### Plasmid-specific genes

The user may add any set of PSGs or remove any of those included with SCAPP.

### Plasmid classification scores

The sequences may be classified using PlasFlow and the PlasFlow classification output file can be provided to SCAPP.

### Algorithm thresholds

Thresholds for finding plasmid gene matches, defining probable plasmid and chromosomal sequences, identifying potential plasmids, filtering them, and many more can all be user-defined. The full software documentation at https://github.com/Shamir-Lab/SCAPP details all of these user options.

## S2 Plasmid-specific genes

We created four sets of plasmid-specific genes (PSGs) by database mining and expert curation:

1. MOB genes: 890 amino acid sequences of plasmid maintenance genes curated by plasmid biologists from the Mizrahi Lab (Ben-Gurion University) and filtered computationally (see details of filtering below).
2. Plasmid ORFs: 4276 nucleotide sequences corresponding to ORFs annotated with ‘mobilization’, ‘conjugation’, ‘partitioning’, ‘toxin-antitoxin’, ‘replication’, or ‘recombination’ from a large set of putative plasmids found by the Mizrahi Lab and then filtered computationally.
3. ACLAME plasmid genes: 4813 nucleotide sequences of genes that make up 96 gene families in the ACLAME database [1] that were manually selected as possibly plasmid-specific. The set of genes was deduplicated and filtered computationally.
4. PLSDB-specific ORFs: 94478 plasmid-specific sequences determined as follows: We used MetaGeneMark [2] to predict genes in the plasmid sequences from PLSDB (v.2018_12_05) [3]. We then counted the number of BLAST matches (> 75% identity match along > 75% of the gene length) to these genes in both PLSDB and bacterial reference genomes from NCBI (downloaded January 9, 2019). We considered each predicted gene that appeared in the plasmids more than 20 times and was > 20 *×* more prevalent in the plasmids than in the genomes to be plasmid-specific.

Sets 1–3 were filtered as follows: We counted matches between the sequences and PLSDB plasmids and NCBI bacterial reference genomes as for the PLSDB-specific ORFs (set 4). We excluded any gene that had more than 4 matches to bacterial genes *and* met one of the following conditions: (1) ≤ 4 matches to plasmid genes and > 4*×* as many matches to bacterial genes as plasmid genes; or, (2) > 4 plasmid gene matches, but ≤ 4*×* as many matches to plasmid genes as to bacterial genes.

## S3 Potential plasmid cycle criteria

Once the set of lightest cycles has been generated, each cycle is evaluated as a potential plasmid based on its structure in the assembly graph, the PSGs it contains, its plasmid score, paired-end read links, and coverage uniformity. A cycle is defined as a potential plasmid if one of the following criteria is met:

1. The cycle is formed by an isolated “compatible” self-loop node *v*, i.e. *len*(*v*) > 1000, *indeg*(*v*) = *outdeg*(*v*) = 1, and at least one of the following conditions holds:
  a. *v* has a high plasmid score *s*(*v*) > 0.9.
  b. *v* has a PSG hit.
  c. *<* 10% of the paired-end reads with a mate on *v* have the other mate on a different node.
2. The cycle is formed by a connected compatible self-loop node *v*, i.e.*len*(*v*) > 1000, *indeg*(*v*) > 1 or *outdeg*(*v*) > 1, and *<* 10% of the paired-end reads with a mate on *v* have the other mate on a different node.
3. The cycle is not formed by a self-loop and has:
  a. Uniform coverage: *CV* (*C*) *<* 0.5, and
  b. Consistent mate-pair links: a node in the cycle is defined as an “off-path dominated” node if the majority of the paired-end reads with one mate on the node have the other mate on a node that is not in the cycle. If less than half the nodes in the cycle are “off-path dominated”, then we consider the mate-pair links to be consistent.

## S4 Simulation of metagenomes with plasmids

To create the simulated metagenomes, we downloaded all completed whole genome bacterial reference sequences from RefSeq (RefSeq database updated on March 11, 2020). We first compiled a list of bacterial strains or species that have been previously identified as prevalent in the human gut from three sources: (1) The list compiled by Alneberg *et al*. [4] (Sup Table 1). (2) Species with abundance > 0.01 in at least one human gut sample from the human microbiome project (HMP1) [5] as estimated by MetaPhLan (abundance table available from https://www.hmpdacc.org/HMSMCP/#data). (3) The “dominant species” identified by Forster *et al*. [6] (Sup Table 5) in the HGG (Human Gastrointestinal Bacteria Genome Collection). We searched for the strains or species on this combined list in the RefSeq database, giving preference to strain level matches. When multiple references appeared (for example, when a listed species has multiple reference strains), we gave preference to those with longer plasmids (>10kbp), followed by those with any plasmid, choosing randomly between references with the same preference. The list of human gut specific bacteria used in the simulations contained 145 references, and is provided in Additional file 2.

For each simulation we first selected from the human gut specific bacteria and then supplemented with randomly selected reference sequences to reach the desired number of genomes. Since the plasmids sequenced with completed whole bacterial genomes are usually long, we also supplemented with a fixed number of shorter (*<*10kbp) plasmids, selected randomly and associated at random with host genomes in the simulation. (5 short plasmids were added in Sim1, 15 in Sim2, 50 in Sim3 and Sim4, 100 in Sim5, 150 in Sim6, and 200 in Sim7.)

Genome abundance and plasmid copy number were assigned using realistic distributions. For genome abundance we used the log-normal distribution (*µ* = 1.5, *σ* = 1), normalized so that the relative abundances sum to 1. This long-tailed distribution mimics the abundance distribution of real microbiome samples. Plasmids were assigned the same abundance as their hosts, and plasmid copy number was assigned according to one of several geometric distributions according to the plasmid length. The parameter of the geometric distribution of a plasmid of length *L* was set to be

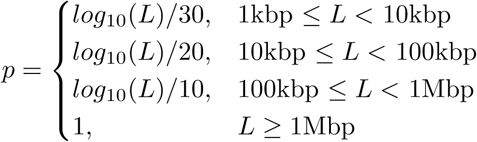

This makes it more likely for shorter plasmids to have higher copy numbers, in accordance with observed plasmid copy number patterns.

Paired-end short reads were simulated from the genome references using InSilicoSeq [7] with the HiSeq error model (default read length = 126bp). To reflect circularity of the plasmids and bacterial genomes, multiple copies of the reference sequence were concatenated before generating reads.

## S5 Experimental settings and evaluation

All metagenomes were assembled using the SPAdes assembler (v3.13) with the --meta option. The default of 16 threads were used, and the maximum memory was set to 750 GB. metaplasmidSPAdes (mpSpades) was run with the same parameters. mpSpades internally chooses the maximal value of *k* to use for the *k*-mer length in the assembly graph. We matched the values of *k* used in SPAdes to these values for each dataset. Defaults were used for all other options for Recycler and SCAPP. In practice, the maximum *k* value was 77 for the simulations and human metagenomic samples, and 127 for the plasmidome and parallel metagenome-plasmidome samples.

For a simulated metagenome, the set of reference plasmids included in the simulation that were covered along > 95% of their length by simulated reads was used as the gold standard. Reads were mapped using BWA [8], and coverage at each base of the reference plasmids was called using bedtools [9].

We used BLAST to match the assembled plasmids to the gold standard plasmid sequences. A plasmid assembled by one of the tools was considered to be a true positive if > 90% of its length was covered by BLAST matches to > 90% of a reference with > 80% sequence identity. The rest of the assembled plasmids were considered to be false positives. Gold standard plasmids that did not have assembled plasmids matching them were considered to be false negatives. Precision was defined as *TP/*(*TP* +*FP*) and recall was defined as *TP/*(*TP* + *FN*), where *TP, FP*, and *FN* were the number of true positive, false positive, and false negative plasmids, respectively. The F1 score was defined as the harmonic mean of precision and recall.

For the human microbiome and plasmidome samples, the set of plasmids serving as the gold standard was selected from PLSDB (v.2018 12 05) [3], a large curated plasmid database. After filtering duplicate plasmids, the PLSDB contains 13469 reference plasmids. The contigs from the metaSPAdes assembly were matched against the plasmids in PLSDB using BLAST. Matches between a contig and a reference plasmid with sequence identity > 85% were marked and a contig was said to match a reference if > 85% of its length was marked. Reference plasmids with > 90% of their lengths covered by marked regions of the matching contigs were used as the gold standard.

The set of plasmids assembled by each method was compared to the gold standard set using BLAST. A predicted plasmid was considered a true positive if there were sequence matches at > 80% identity between the plasmid and a gold standard plasmid that covered more than 90% of their lengths.

Note that in the case of the real samples, if two assembled plasmids matched to the same reference gold standard plasmid sequence(s), then one of them was considered to be a false positive. This strict definition penalized methods for unnecessarily splitting potential plasmid genomes into multiple different plasmids. If there were multiple gold standard reference plasmids that were matched to a single assembled plasmid, then none of them was considered as a false negative. The precision, recall, and F1 score were calculated as for the simulation.

**Table 1.** Full performance on simulated metagenome datasets. The gold standard is the number of plasmids in the simulation that are covered by simulated reads (# covered).

| Sample | # covered | Recycler |  |  | mpSpades |  |  | SCAPP |  |  |
| --- | --- | --- | --- | --- | --- | --- | --- | --- | --- | --- |
|  |  | precision | recall | F1 | precision | recall | F1 | precision | recall | F1 |
| Sim1 | 9 | 57.1 | 44.4 | 50.0 | 100 | 11.1 | 20.0 | 80.0 | 44.4 | 57.1 |
| Sim2 | 37 | 60 | 32.4 | 40.1 | 100 | 24.3 | 39.1 | 56.5 | 35.1 | 43.3 |
| Sim3 | 136 | 52.5 | 23.9 | 32.8 | 96.3 | 19.4 | 32.3 | 81.3 | 29.1 | 42.9 |
| Sim4 | 132 | 62.9 | 30.2 | 40.8 | 100 | 22.3 | 36.5 | 86.3 | 34.1 | 48.9 |
| Sim5 | 253 | 55.6 | 25.7 | 35.2 | 96.2 | 20.5 | 33.8 | 77.2 | 34.3 | 47.5 |
| Sim6 | 368 | 51.4 | 19.7 | 28.5 | 96.6 | 15.8 | 27.1 | 72.9 | 24.4 | 36.5 |
| Sim7 | 410 | 62.9 | 20.6 | 31.1 | 95.7 | 16.5 | 28.1 | 75.9 | 27.6 | 40.5 |

**Figure S1.**
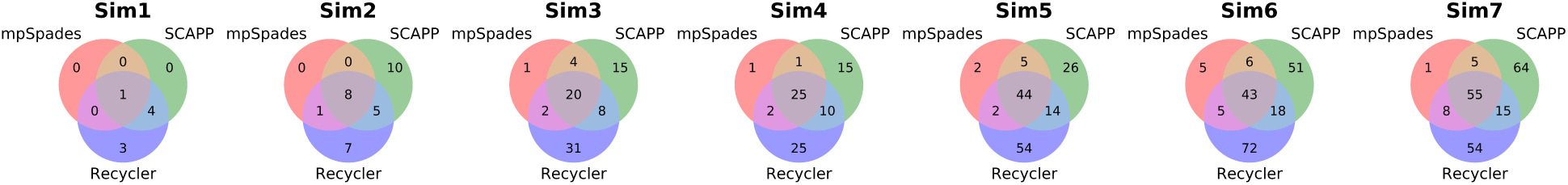
Plasmid overlap between tools in simulation. Overlap of the plasmids assembled by the tools on each of the simulated metagenomes.

For the parallel metagenome-plasmidome sample, plasmidomic reads were aligned to the plasmid sequences and metagenome assembly contigs using BWA [8]. Coverage at each base of each metagenomic contig was called using bedtools [9].

To compare the overlap between plasmids identified by the different tools, we considered two plasmids to be the same if their sequences matched at > 80% identity across > 90% of their length. For visualization purposes, when two plasmids in one tool match one plasmid in another, they are represented as one overlap in the venn diagram (Figures S1 and S3).

## S6 Extended results for simulated datasets

Table 1 reports the full precision, recall, and F1 performance results for all tools on the simulated metagenome datasets. Figure S1 shows the overlap between the plasmids assembled by each tool in the simulated metagenomes.

## S7 Extended results for human metagenomes

Figure S2 presents the F1 scores of the plasmid assemblers across all human gut metagenome samples. Table 2 reports the full results and the number of plasmids assembled by each tool and the median plasmid length for each of the human gut microbiome samples.

**Figure S2.**
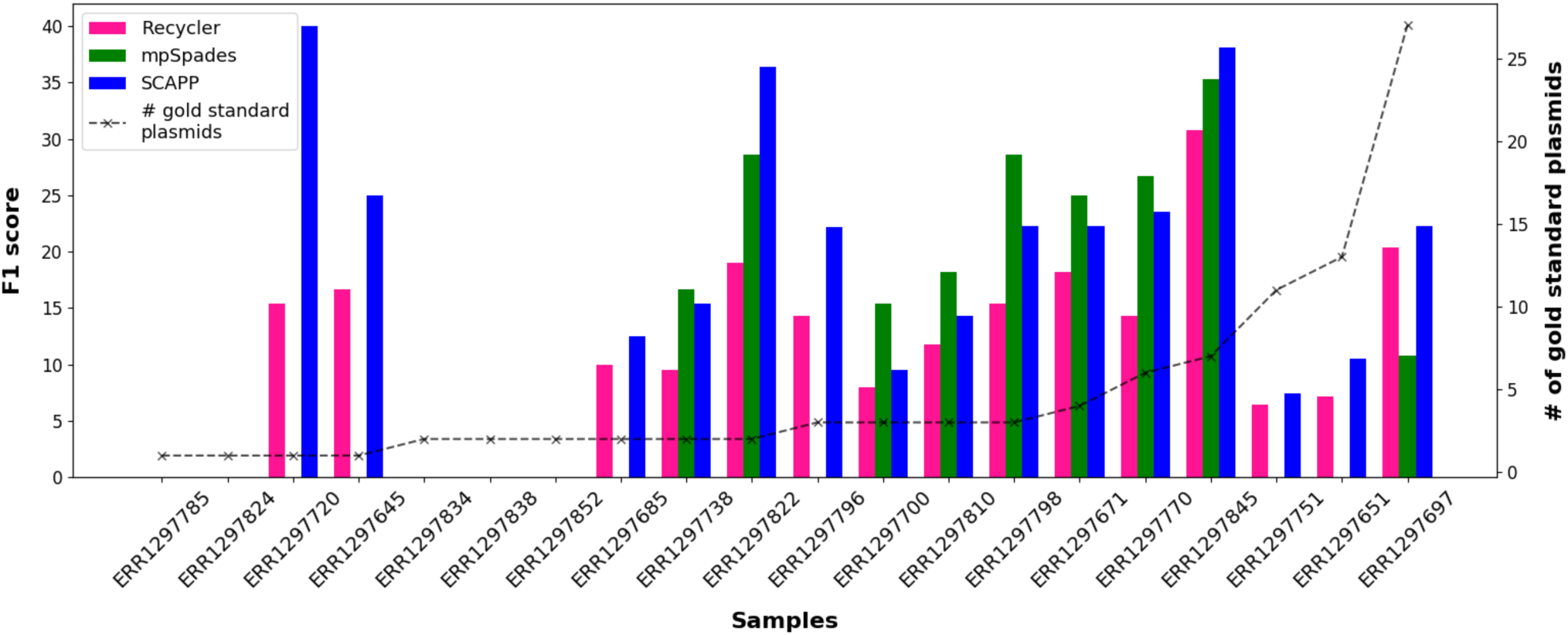
Results on 20 human gut metagenomes. F1 scores of the plasmids assembled by Recycler, mpSpades and SCAPP in the human gut microbiome samples (accessions given on x-axis), calculated using PLSDB plasmids as the gold standard. The dashed line shows the number of gold standard plasmids in each sample. Where bars are omitted the F1 score was 0.

**Table 2.**
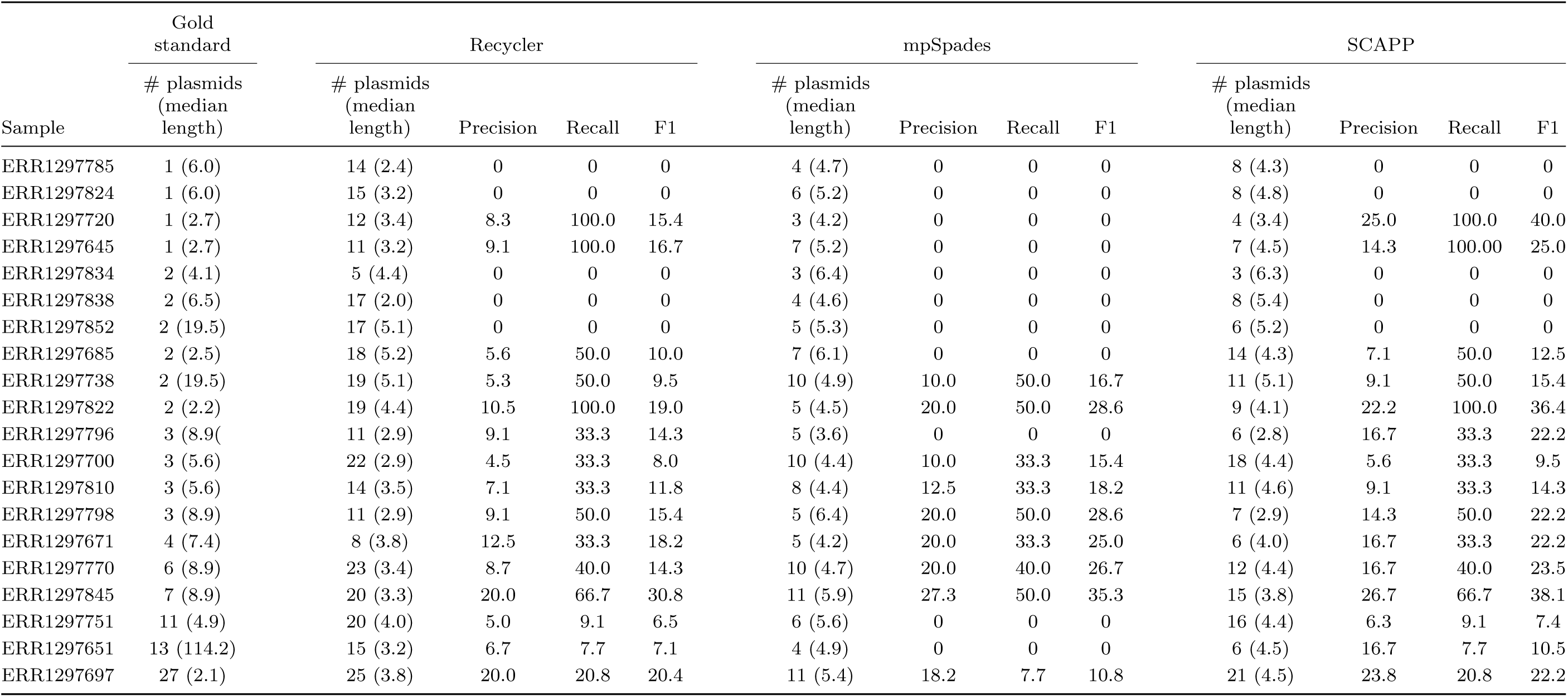
Full results on the human gut microbiome samples and number of plasmids and median lengths (in kbp).

**Figure S3.**
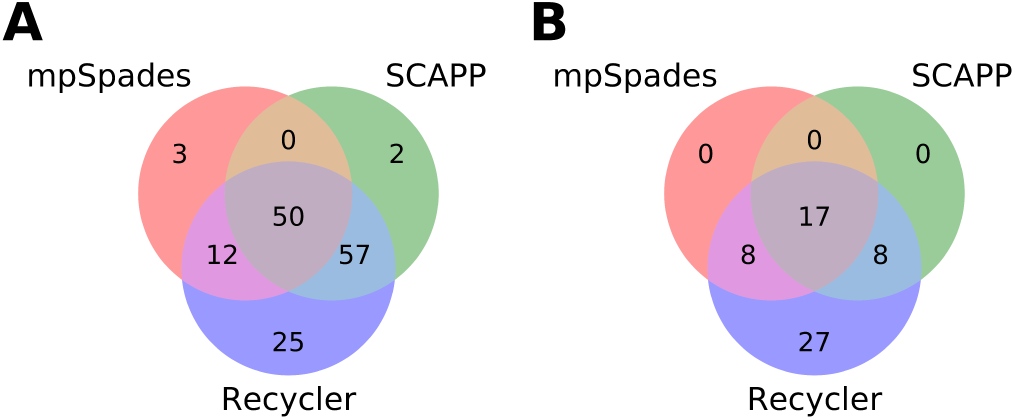
Number of plasmids assembled by each tool on the parallel samples. A: Plasmidome sample. B: Metagenome sample. Discrepancies between the numbers in the diagram and Table 4 are due to cases of overlaps between two plasmids in one tool to one plasmid in another, which were counted as one.

## S8 Extended results for parallel plasmidome-metagenome

Figure S3 shows the overlap between the plasmids assembled by the tools in the parallel cow rumen plasmidome and metagenome samples.

Figure S4 shows the annotations of the gene functions and hosts for the plasmids assembled in the rumen plasmidome.

**Figure S4.**
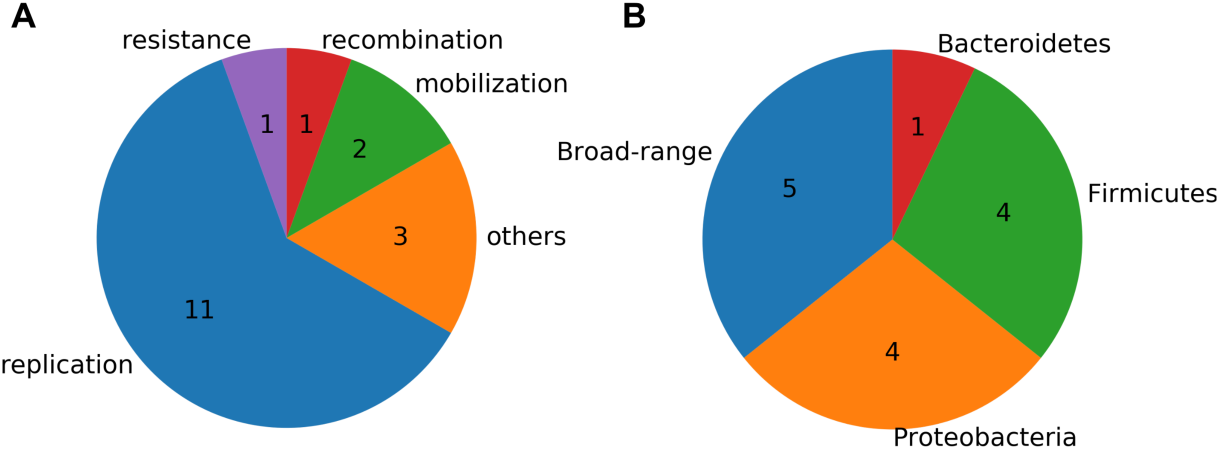
Annotation of genes on the plasmids identified by SCAPP in the rumen plasmidome sample. A: Functional annotations of the plasmid genes. B: Host annotations of the plasmid genes.

## Notes

### Competing Interest Statement

The authors have declared no competing interest.

### Summary of Updates

Revised draft - additional simulations and analysis.

